# The limbic circuit for response vigour

**DOI:** 10.1101/600973

**Authors:** Bernd Porr, Pedro Gomes, Thor Andreassen

## Abstract

Motivation can be defined as the amount of energy an animal puts into actions to achieve goals. This not a static property but can be learned. Recently, abstract models have been developed for this kind of learning which propose that tonic dopamine is responsible to energise behaviour. However the underlying circuitry is still not understood and it is still unclear how tonic dopamine acts on the motor system to energise behaviour. In this paper we present a limbic system model of response invigoration which is based on neurophysiological data and anatomical connections. We argue that the indirect pathway from NAcc core D2 neurons via the ventral pallidum to the SNr conveys response vigour, while the direct pathway from NAcc core D1 neurons is responsible for action selection. Once this detailed anatomically mapped circuit has been established we then condense this realistic model into an abstract model which then shows that tonic DA acts in a multiplicative way on the motor output and thus is able to energise behaviour. We demonstrate the model with both a simulated and real robot experiment.

## 1 Introduction

The mesolimbic dopamine system has long been implicated in the mediation of reward-related and goal-directed behaviours (Wise, 2004a). There are widely considered to be two main modes of dopamine release, tonic and phasic. The phasic dopamine signals, the result of the bursting activity of dopamine neurons, have been implicated in reinforcement learning which has been widely accepted coding a reward prediction error (Dayan, 2002; Schultz et al., 1997; Sutton and Barto, 1987, 1998). In this paper we focus on the slower changing levels of tonic dopamine which have been related to motivation in the sense of energising behaviour (Cofer, 1981). In terms of reinforcement learning (Niv, 2007) suggests that the level of tonic dopamine is related to the average rate of rewards being received by an animal. It is proposed that it is beneficial for an animal to become more motivated, expend more energy and to work faster in an environment where there are more rewards available.

Not only do animals show invigoration in response to reward delivery, animals also become invigorated in response to anticipation of reward delivery. The level of tonic dopamine would be expected to rise when the animal is placed in an environment where rewards have previously been received or when the animal is presented with a stimulus predicting the availability of a reward. Indeed the existence of an ‘anticipation-invigoration mechanism’ which allows the presentation of a conditioned stimulus to bring about an increase in response vigour was originally proposed by Cofer and Appley (1964).

While the role of dopamine as a neuromodulator has been widely investigated (Kilpatrick et al., 2000; Mirenowicz and Schultz, 1994; Schultz et al., 1997), the underlying network generating this activity is still poorly understood. In this paper we will investigate the network controlling tonic dopamine release and how it impacts on motivation and thus energises behaviour. In particular we propose that dopamine influences different pathways in opposing ways mediated by the differential actions of dopaminergic receptors in the Nucleus Accumbens (NAcc). We will present a computational model of the areas of the limbic system we believe are involved in the release of mesolimbic dopamine and behaviour selection where the level of tonic dopamine represents the average reward rate in a food retrieval task and affects the response rate of the agent accordingly.

The outline of the paper is: first, we describe the different nuclei which are responsible for conveying response vigour and motivation. Then, we will describe the signal flow between these nuclei, derive a mathematical model which is closely mapped on the anatomical structure, derive an equivalent abstract model and then as the final step implement it in two behavioural setups whereas the one is a simulated and the other a real robot.

### 1.1 The limbic circuit conveying motivation

In this section we are going to present the brain circuitry which we propose to be responsible for controlling response vigour and motivation. Our main goal is here to show how tonic dopamine differentially influences activity down to the motor output stages which involves chains projecting from one nucleus to the next one which are often inhibitory. This results in chains of nuclei which are inhibited, disinhibited, inhibited and so on. At the end of this section we will then conclude which pathway is inhibited by tonic DA and which one is excited. The change of balance between inhibition and disinhibition will then change the motor activity and in turn response vigour.

Fig. 1 shows the relevant limbic areas of interest which are: orbitofronal cortex (OFC), medial pre-frontal cortex (m-PFC), the nucleus accumbens with its subdivistions core/shell, ventral pallidum (dl-VP, m-VP), lateral hypothalamus (LH), substantia nigra reticula (SNr) and the ventral tegmental area (VTA). In particular, we propose that tonic dopamine has a differential influence on the different core neuron subpopulations called “core-D1” and “core-D2” which carry the D1 and D2 dopamine receptors respectively. These in turn project directly and indirectly to the motor output of our model, the SNr. We are now going to describe these different limbic areas in greater detail.

**Figure 1.**
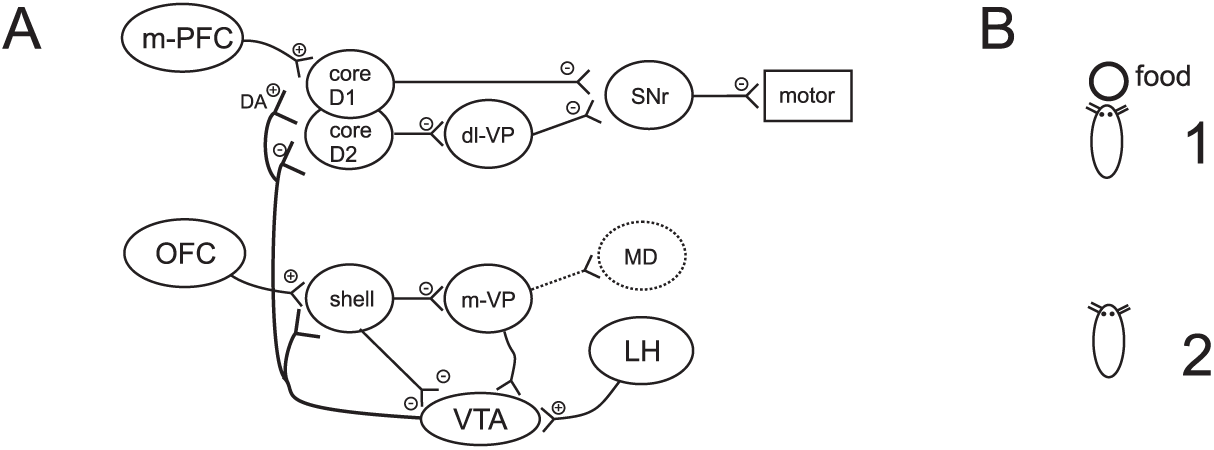
The tonic dopamine vigour model based on anatomical data. (A) and experimental setup (B). Abbreviations: m-PFC - medial prefrontal cortex, OFC - orbitofrontal cortex, dl-VP, m-VP - dorsolateral and medial ventral palladium, VTA - ventral tegmental area, LH - lateral hypothalamus, MD - mediodorsal thalamus, SNr - substantia nigra pars reticular, core D1/D2 - nucleus accumbens core neurons with dopamine D1/D2 receptors, shell - nucleus accumbens shell region

#### 1.1.1 Nucleus Accumbens (NAcc)

The nucleus accumbens has been described as the ‘limbic motor interface’, proposed to be the final structure involved in the initiation of movement before the information is relayed to the basal ganglia where lower level planning can take place. Its role is to integrate motivational and emotional information from other limbic and cortical structures (Mogenson et al., 1980).

There are two main subdivisions of the NAc, the core and the shell, which can be distinguished by their unique connectivity (Sesack and Grace, 2010). The NAc core receives inputs from the dorsal-medial prefrontal cortex and the hippocampus while the NAc shell receives input from the basolateral amygdala, the orbitofrontal cortex and the hippocampus (Brog et al., 1993). The shell is rather involved in motivational aspects, the core as being closer to the dorsal striatum is responsible for motor initiation and feeding motor signals to more dorsal decision making circuits via cortico striatal loops (Haber, 2003).

Here, the pathways to the motor circuits are of special importance to our motivation model. There are two distinct output pathways from the NAcc core which have its origins from the two subpopulations of neurons in the NAcc core: the one subpopulation carries mainly D1 receptors and the other one carries mainly D2 receptors (Humphries and Prescott, 2010; Kelley, 2004). The D1 receptor carrying neurons feed directly into the SNr and are able to *inhibit* tonically active SNr neurons, thus the NAcc core is able to *disinhibit* motor programs. We call this pathway in accordance to the basal ganglia terminology the *direct pathway*. On the other hand there is an *indirect* pathway via the VP to the SNr originating from the NAcc core. In contrast to the *direct* pathway these neurons in the indirect pathway carry mainly D2 receptors which are inhibitory in nature. D2 receptors are very sensitive to low DA concentrations (Grace, 2000) and will react to the tonic DA concentrations. This indirect pathway will play a central role in our response vigour model and we will come back to it once we have introduced the other nuclei.

#### 1.1.2 Ventral pallidum (VP)

The ventral pallidum contains mainly GABAergic neurons which are tonically active (Floresco et al., 2003) and their efferents target a variety of subcortical structures (Groenewegen et al., 1993). This means that the VP inhibits its target regions while not being stimulated.

Two pathways are routed through the VP have distinct roles in our model. The first distinct pathway is the aforementioned one responsible for motor initiation from the core/D2 subpopulation which projects to the dl-VP. Because the dl-VP neurons are tonically active they inhibit the SNr and other target areas. Coming back to our tonic DA model, a rise in tonic DA will inhibit the core D2 subpopulation which in turn will disinhibit the dl-VP neurons.

The other pathway runs from the shell to the m-VP then to the VTA and exerts a strong inhibitory influence on the VTA due to the tonically active neurons in the m-VP (Floresco et al., 2003). Taking into account that the NAc shell inhibits the m-VP, an activation of the NAc shell then disihibits the VTA which leads us to the VTA and its release of dopamine. This pathway will be crucial to control the level of tonic DA.

#### 1.1.3 Substantia Nigra pars reticular (SNr)

The SNr is the output stage in our model and is responsible for the activation of motor programs (Kelley, 2004). As for the VP the SNr has tonically active neurons which ultimately inhibit motor programs via the standard basal ganglia pathways (Groenewegen et al., 1999; Nicola, 2007). Recall that both the NAcc core and the dl-VP project to the SNr and are therefore able to inhibit SNr neurons which then in turn disinhibits motor programs.

#### 1.1.4 Ventral Tegmental area (VTA)

The ventral tegmental area has a high density of dopaminergic neurons which have two modes of operation: tonic activity and phasic activity. While the circuitry generating phasic activity has been extensively investigated in the light of reinforcement learning (Wörgötter and Porr, 2005) we focus on the circuitry around tonic DA.

Tonic dopamine is released due to the fact that, under normal circumstances, around 50% of midbrain dopamine neurons are spontaneously active, firing irregularly with a rate of between 2 and 10 Hz (Grace et al., 2007). Factors which alter the number of these tonically active neurons are thought to contribute to the level of tonic dopamine in the extrasynaptic space (Grace, 2000; Grace et al., 2007). Due to its considerably lower concentration (by a factor in the order of 100,000 - (Grace, 2000)), tonic dopamine is thought to exert its effects primarily through the high affinity D2 class dopamine receptors (Richfield et al., 1989; Schultz, 2007) which leads us to the target regions of the VTA.

While the VTA has a whole host of target regions, of special importance is the projection from the VTA to the NAc. As mentioned in section 1.1.1 about the NAcc core, tonic DA can stimulate D2 receptors in the NAc which in turn will inhibit those NAc neurons carrying a high number of D2 receptors. We propose that the response vigour is controlled by tonic DA acting on the D2 receptors, especially in the NAcc core. The higher the tonic activation the less D2 carrying NAcc core neurons become active which makes it easier downstream to execute actions. We will explain now in greater detail how tonic DA influences the decision making network.

#### 1.1.5 Tonic DA and action invigoration

We are now going to describe in detail how actions are energised by tonic DA. As mentioned before there are two distinct pathways for action selection from the NAcc core to the SNr which is our motor output: the *direct* pathway from the core to the SNr and the *indirect* pathway via the VP. We propose that tonic DA influences the response vigour via the indirect pathway while the direct pathway performs the actual action selection.

Let us first describe the direct pathway: it is called direct because the subpopulation of NAcc core neurons carrying D1 receptors inhibit the SNr directly (Humphries and Prescott, 2010) which then in turn disinhibits motor programs. The acquisition of these motor programs has been extensively described in (Thompson et al., 2010a). Also in our model these motor programs are learned and initiated via this pathway.

Central to our response vigour model is the indirect pathway from the NAcc core neurons which carry mainly D2 receptors. We propose that this pathway energises motor programs and thus controls how vigorously they are executed. This can be seen if we look now at the overall effect tonic DA has on action activation and we count now how many subsequent inhibitions (-) we accumulate to assess the overall result: is it finally inhibition or disinhibition? A rise in tonic DA stimulates D2 receptors in the D2 carrying population of the NAcc core which inhibit (-) the NAcc core D2 carrying neurons. These NAcc core neurons inhibit the dl-VP (-) which in turn disinhibits the SNr (-) which has an inhibitory (-) influence on motor programs. In total we have therefore (-,-,-,-) a disinhibitory effect on motor programs when tonic DA rises. This means that tonic DA energises behaviour via the indirect pathway.

After having described the anatomical connections and provided a qualitative model we now need to turn this into a mathematical model so that we can model animal behaviour in both a simulation and a robot.

## 2 Methods

The previous section has provided the anatomical structure and also given a qualitative description of the model. In this section we are now going to describe its function and formalise it. We will first map the anatomy directly onto equations one by one. At the end we will show that these can be condensed into one equation for the response vigour where tonic DA energises behaviour, essentially acting as a multiplicative factor.

### 2.1 Approach behaviour

Before we can talk about the motivational aspects of learning we first need to describe briefly how a naive animal learns to approach food from the distance which has been covered extensively before (Prescott et al., 2006; Thompson et al., 2010b). For instructional purposes we use a simple behavioural thought-experiment (see Fig. 1B). Later we will use a similar experiment for the simulation and the robot where the only difference will be practical issues such as sensor and motor processing. In this section we first focus just on the learning of the motor task and then further down describe how this can be energised.

The task of the animal is to learn to see and then approach the food from the distance. This is achieved by correlating a proximal sensor signal with a distal one. The actual computation is achieved in the core, in particular in the NAc core D1 subpopulation, Input to the NAc core neurons with a high D1 receptor density is modelled as originating from the dorsal medial prefrontal cortex (dmPFC). The dmPFC is implicated in working memory (Compte, 2006; Seamans et al., 2003) and is considered, from the perspective of this model, to hold representations of relevant stimuli such as the position in the sense of a “where” signal. The dmPFC is modelled as relaying two signals to the NAc core, one distal signal (Fig 1B2) which represents a conditional stimulus (CS) and one proximal signal (Fig 1B1) which represents the unconditional stimulus (US). The distal signal is activated when a stimulus is within sight of the modelled agent or robot and is representative of the distance to the stimulus. The proximal signal is activated at close proximity to the stimulus and represents a natural curiosity in the object, allowing the agent to make a final approach when it otherwise might not. Again, US and CS are modeled as simple step functions indicating if an object is close by (US) or seen from the distance (CS). Both functions are filtered by lowpass filters *h*_*CS*_(*t*), *h*_*US*_(*t*) and acting here as memory traces (see the appendix for the parameters).

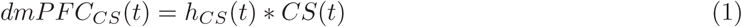

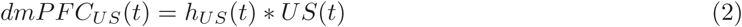

The * from now on denotes a convolution (i.e. filtering) operation.

Activity within the NAc core D1 subpopulation is calculated as the sum of the weighted *dmPFC*_*CS*_(*t*) and unweighted *dmPFC*_*US*_(*t*) afferents:

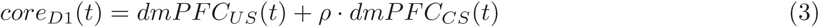

where the activity of the core D1 is composed of a fixed curiosity reaction via *dmPFC*_*US*_(*t*) and a learned approach action via ·*dmPFC*_*CS*_(*t*) where *ρ* is the weight determining how strong the agent acts once it has seen an object from the distance. If there were more landmarks then more inputs and weights could have been added (Thompson et al., 2010a).

As a next step we need to describe how the core learns the approach behaviour. We start with the naive animal which is wandering around in the arena (Fig 1B). Eventually the animal will reach a pot filled with food by chance (Fig 1B1). This triggers the primary reward system, the lateral hypothalamus (LH). Neurons in the LH are known to be active at the point (or just after) reward delivery and the area has previously been modelled as a ‘primary reward’ region (Brown et al., 1999). Reward input (*reward*(*t*)) to the model thus occurs through the LH, where the resulting activity is modelled as a lowpass filter response *h*_*reward*_ from an impulse reward input *reward*(*t*) and is given by the following equation:

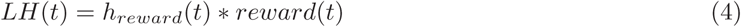

Learning is now established via the LH-VTA pathway. The LH has a strong excitatory influence on the ventral tegmental area (VTA - bottom of figure 1) and is able to cause phasic spiking in VTA neurons (Grace et al., 2007) which we call *burst*(*t*) whenever the LH is activated. We will provide the exact equation later once we have introduced the complete circuitry around the VTA. For now we just need to know that the VTA burst switches on learning in the Nacc core and in turn associates the stimulus representing the food pot with an approach behaviour (Porr and Wörgötter, 2007). Bursts of phasic dopamine are known to activate D1 receptors in the NAc, which, along with glutamate release contributes to the induction of long term potentiation (LTP) of NAc synapses (Schotanus and Chergui, 2008). Thus learning in the core allows for an appropriate approach behaviour to be engaged on the presentation of a rewarded stimulus from the distance (Fig 1B2).

More formally the phasic, or burst mode of dopamine action can be interpreted as a third factor in a three factor learning paradigm (Porr and Wörgötter, 2007) implemented in the NAc which essentially switches on (differential) Hebbian learning at the moment of the reward. The change of the weight *ρ* is modelled using the three factor learning rule, ISO3 (Porr and Wörgötter, 2007), where the third factor is provided by the dopamine burst and the core D1 weight *ρ* is then updated according to the following equation:

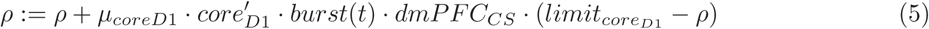

which has as a first factor the derivative of the output of the core, as a second factor the DA bursting activity and as a third the input from the dmPFC. The last term 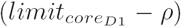 prevents the weight from growing endlessly and saturates it at the soft limit of *limit*_*core*_.

After a few trials the animal should be able to approach the food pot from the distance by channeling the appropriate environmental stimulus (landmark, smell, etc) via the dmPFC through the NAc core to the motor outputs. Once the animal has learned this association it will approach the food pot instantly. The next section describes the tonic dopamine which modulates how vigorously the animal approaches the food pot.

### 2.2 Response vigour

After having learned to approach food from the distance we now describe our model for the response vigour, namely how vigorously an animal approaches a landmark which has been associated with a reward (see Fig. 1B2). Again, we start with the naive animal which just wanders around in its environment. As previously described, encountering the primary reward (see Fig. 1B1) triggers a burst in the LH which in turn switches on learning but this time we focus on the NAc shell. Learning in the NAc shell establishes the association between the reward and the distal stimulus which in turn controls the response vigour via tonic DA. This is in contrast to the NAc core which establishes stimulus - action associations.

In order to understand how the shell manages to control the response vigour we need to look into its afferents which originate primarily from the orbital frontal cortex (OFC) (Thompson et al., 2010b; Zahm, 2000). Neurons in the OFC respond to stimuli from the environment and as such are modelled as conveying information about active stimuli to the NAc shell. The OFC is shown on the left hand side of figure 1A. Activity along the pathways connecting the OFC to the NAc is defined as the lowpass filtered response of signals from the surrounding environment providing memory traces of these signals:

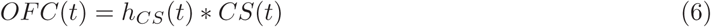

The activity of the shell is computed as a weighted (learned) input from the OFC and an unweighted input from the LH (Thompson et al., 2010b):

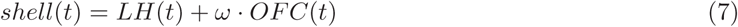

where *ω* is the weight associating the OFC activity with a reward.

Again, learning itself is established by phasic DA activity at the moment of encountering a primary reward and this triggers learning in the shell:

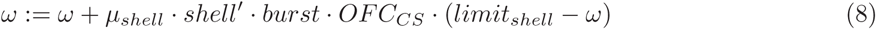

At moment of a reward there will be activity in the shell due to its direct input from the LH, a burst of DA activity and, if a CS has been present, also input from the OFC so that the product of these three factors will increase the weight *ω*.

Having the shell activity we can now finally formulate the overall activity in the VTA:

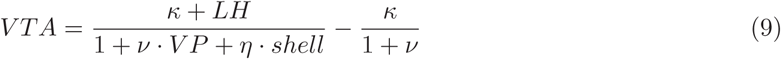

where *v >> η* which indicates a strong influence of the VP on the VTA and a weaker input from the NAc shell to the VTA. The constant *κ* defines the tonic activity of the DA neurons which can then be suppressed by a strong input from the VP and a weak one from the shell. The strong input from the VP is of special importance to our model as it controls the tonic activity of the VTA neurons. The bursting activity at the moment of the reward delivery is generated by the LH input. It is know that this DA burst becomes smaller and this is achieved by the direct pathway from the shell to the VTA via the weight *η*.

The bursting activity of the VTA is simply the rectified highpass filtered version of DA and now be formulated:

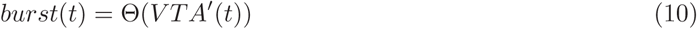

This generates a short spike when the activity in the VTA rises which can be causes by a sudden input from the LH or also by the rise of DA through disinhibition (2nd order conditioning, see (Thompson et al., 2010b)).

The DA tonic dopamine signal is detected by simply lowpass filtering the overall VTA activity:

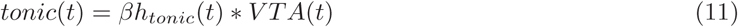

with the lowpass filter function *h*_*tonic*_(*t*) and *β* = 1. In the case of the D2 antagonist intervention we set the tonic gain factor *β* = 0.1.

We now need to describe how tonic DA influences the decision circuitry which is ultimately the SNr which disinhibits motor programs. We propose that this is achieved by stimulation of the core D2 population which then in turn affects the SNr via the indirect pathway. We explain this now in a formal way.

The D2 receptors on the NAcc core subpopulation are inhibited by tonic DA. We assume that their activity is constant in this experiment because it is not altered during the acquisition but rather in scenarios where behaviour needs to be inhibited (Humphries and Prescott, 2010) which is not the subject of this paper:

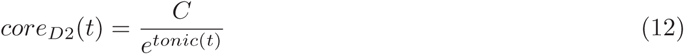

where *C* is a constant. For the shunting inhibition we use the exponential function in the denominator of Eq. 12 instead of 1 + *x*. We choose the exponential here instead of 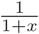 because through our chain of inhibtions and disinhibitions this would lead to unnecessary complex terms and would just obliterate the final result (and note that *e*^*x*^ ≈ 1 + *x*). The exponential will allow us even after several inversions to see if the final result is inhibition or disinhibition and will lead to a compact result.

The output of the core D2 subpopulation then projects to the dl-VP:

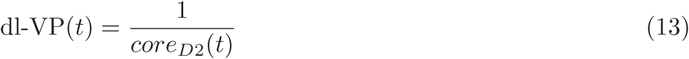

Taking into account both the direct and indirect pathway, the one arriving from the core D1 subpopulation (Eq. 3) and the other one from the dl-VP (Eq. 13) we have an overall SNr activity of the form:

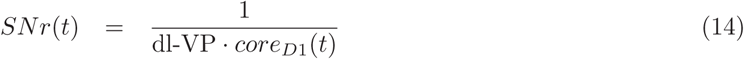

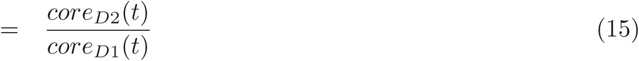

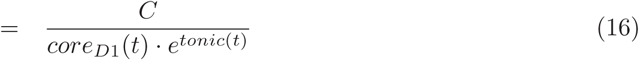

which means that activity in the NAcc core D1 subpopulation is able to inhibit SNr activity and that tonic DA activity can do the same. Recall that the SNr exhibits high tonic activity which inhibits downstream motor programs. This means that when the activity of certain SNr neurons drops then certain motor programs are activated or energised. The vigour of a motor program is coded by the reduction of inhibition from the SNr. Thus, the less the SNr is active the stronger a motor program is executed. To guarantee that no motor action is executed we introduce a threshold which needs to be exceeded so that a motor program can be executed. The actual motor activity is then calculated by inverting Eq. 16 and introducing a threshold.

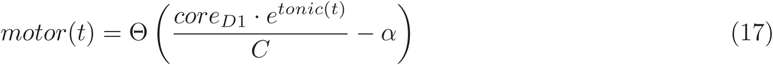

where *α* is a threshold so that at rest no action is triggered.

Overall we have shown that the tonic activity has a multiplicative effect on the actions initiated by the NAcc core neurons. The more tonic DA is present the stronger the response vigour and also the ability to initiate an action because it will be easier to overcome the threshold.

Eq. 17 is our *abstract model* of the reponse vigour which we have derived from our biologically realistic model shown in Fig. 1. As a circuit diagram the abstract model is shown in Fig. 2. Preprocessed CS and US information is relayed from the dmPFC into the NAc core. Learning takes place as per the three factor learning rule shown, where the phasic dopamine signal provides the third factor to switch on learning. The tonic dopamine signal which codes the response vigour then has a multiplicative effect on the output of the learner which is then sent over a constant threshold. We are going to use this model in our simulations and robot experiments below.

**Figure 2.**
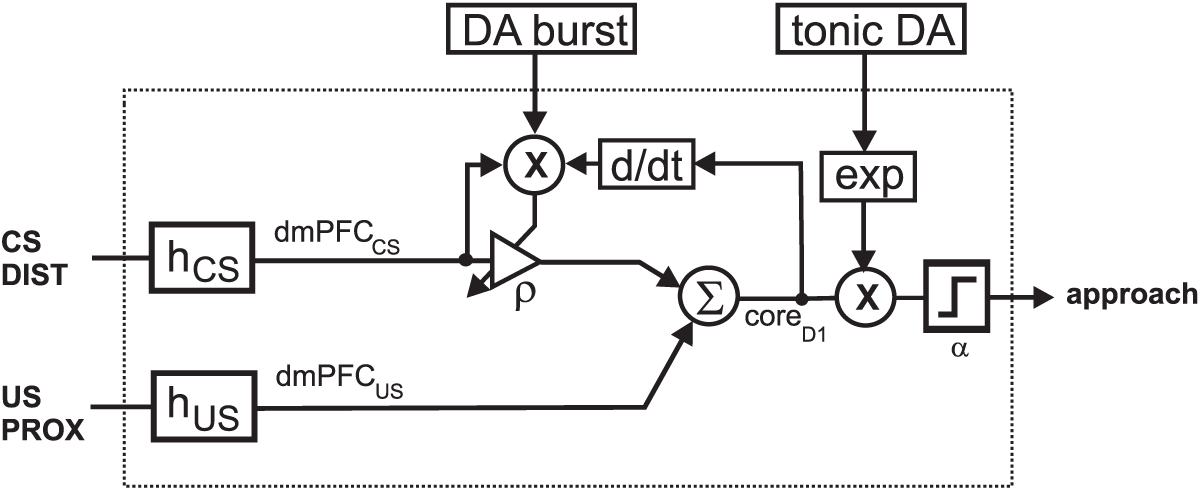
The *abstract model* of response vigour. “CS DIST” and “US PROX” represent the proximal and distal signals in the environemt which can be interpreted as US and CS. These signals are then smoothed by *h*_*C*_*S* and *h*_*U*_ *S*. × represents a multiplication with three factors: DA burst, filtered distal signal *dmPFC*_*CS*_ and the derivative of the output *core*_*D*1_. The product changes the weight *ρ*. The output *core*_*D*1_ is multiplied with the exponential of tonic DA and then sent over a threshold with value *α*. The output of the threshold then enables the motor behaviour.

In summary we have shown that the response vigour is conveyed to the motor circuits via the “indirect” pathway of the limbic system by stimulating the Nacc core subpopulation which carries D2 receptors with tonic DA which then projects to the VP and then from there to the SNr which has an overall multiplicative effect on the motor activity and thus engergises the selected actions.

### 2.3 Experiments

Our methods to test this model are both simulation and real robot runs which are based on the runway task conducted by Marx and Brownstein (1963). In the original experiment, the progress of rats along a 14 ft by 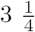 in was timed (see Fig 3). Timing began when a start door was opened and ended when the animal entered an end box containing a food reward. Along with the total running time, the experimenters timed the forward progress (FP) of the rat. FP time only included the time that the rat spent making progress toward the end box (see the two dotted lines in Fig 3A). If the rat stopped or turned around, the timer was stopped and only started when the rat again reached its position of furthest progress.

**Figure 3.**
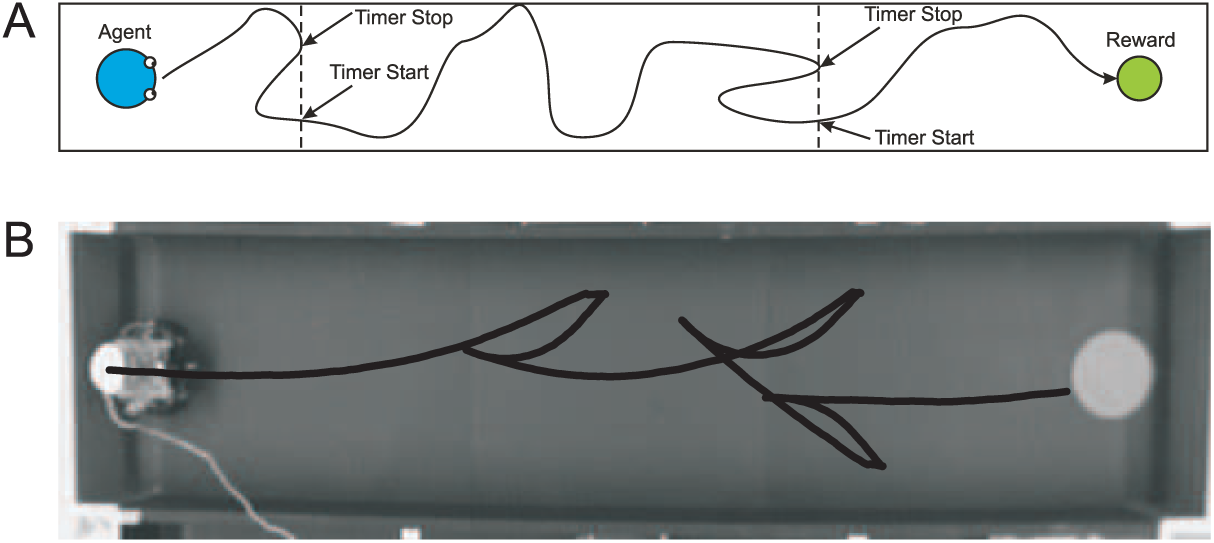
Illustration of the runway task used to demonstrate the tonic dopamine vigour model (Marx and Brownstein, 1963). A shows the simulation enviornment and B the real robot. The task of the agent is to find the reward on the right by moving down the alley from left to right. In both cases an actual trajectory before learning is shown. As in the paper of (Marx and Brownstein, 1963) only the forward speed towards the reward is measured.

To test that tonic dopamine indeed has the effect on response vigour we applied a D2 antagonist into the NAcc shell which should reduce the drive of the agent substantially because it would prevent the tonic dopamine inhibiting the Nacc D2 subpopulation which then dishibits the SNr via the indirect pathway.

#### 2.3.1 Simulation

The simulation provided a free running environment where the agent could engage in basic exploratory behaviour and then eventually would find food at the end of the runway. The run had a width of 1400 pixels and a height of 100 pixels with a marker which triggered the reward input to the learning model place at one end (see again Figure 3A).

In the original experiment by Marx and Brownstein (1963), only one run was made per rat per day. To simulate this, the model was allowed to run between each trial without input until the internal representation of tonic dopamine had settled back to baseline levels.

#### 2.3.2 Real Robot

The driving robot scenario used a run of length 15 ft and width 1.15 ft. A photograph of the physical experimental setup used is shown in Fig. 3B. The reward input to the learning model was triggered when the robot reached the red circular marker at the finishing end of the run.

The robot was an adapted Rug Warrior (www.robotbooks.com/rug_warrior.htm) with a front mounted camera for navigation and two bumper switches for collision detection. A simple thresholding technique was applied in real time to each frame retrieved from the camera to determine which (if any) pixels within the image were part of the marker. An average of the coordinates of the matching pixels was then used to calculate the position of the marker within the image. The height of marker on the image was used to calculate an approximation of the distance to the marker and used as the distal input to the learning model. If the value of the distal input was above 0.9, the proximal input was also triggered. If the output of the learning model determined that the robot should approach the marker, a steering correction was calculated from the horizontal position of the marker within the image so that the robot drove toward the marker.

During exploratory behaviour the robot made small, random changes to its trajectory. Collisions with the side of the run were detected through the front mounted bumper switches. On colliding with the wall, the robot reversed and performed a randomised turn such that the resulting trajectory was arbitrary, but within 90 °of the direct approach angle of the marker.

Each individual trial was filmed using an overhead camera and a tracking algorithm applied to the footage in order to determine the forward progress time. The same pharmaceutical interventions were tested as in the robot scenario as with the simulation.

## 3 Results

We structure this section similar to the previous one that we first present the results for the simulation and then the ones for the real robot run.

### 3.1 Simulation

Similar to the original experiment, the average forward progress speed of the agent was calculated from the recorded forward progress times for each trial. These are plotted in Fig. 4A and Fig. 3A shows actually a trajectory from one of these runs. On the x axis we have the number of trials which represent one approach behaviour down the runway. After 72 trials the system was reset and the simulated agent had to learn again. This was repeated 20 times and then the average and standard deviation was calculated.

**Figure 4.**
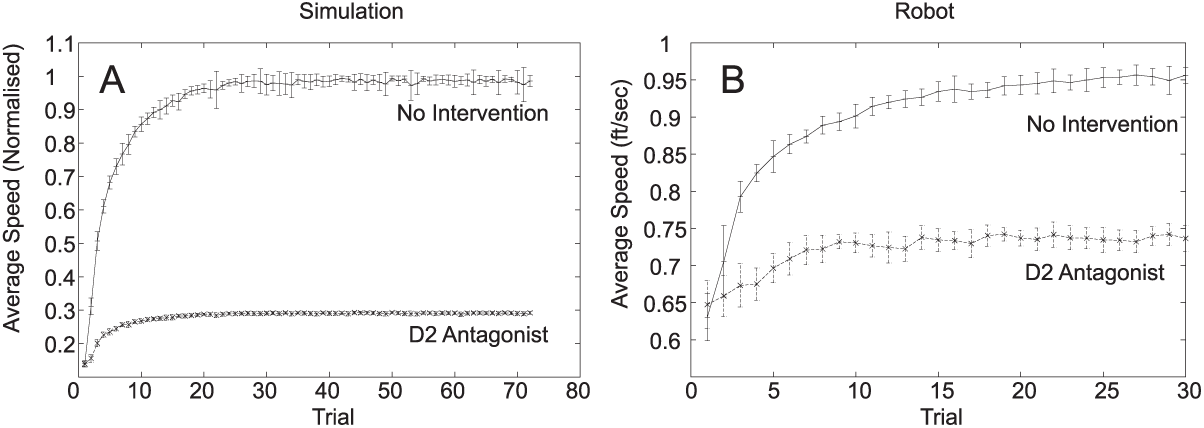
Results from the simulation (A) and the robot experiment (B) Both, the simulation and the robot experiment were run with an optimally working limbic system “no intervention” and under the influence of an dopamine D2 receptor antagonist “D2 antagonist”. The x axis shows the number of trials where every trial is one run down the alley. The y axis shows the average speed of the robot where in the simulation this is normalised to the final speed of the robot in the control scenario “no intervention”. In the robot experiment the the y axis shows the actual speed of the robot.

In the case of no pharmaceutical intervention, the agent shows a progressive improvement in average speed over each consecutive trial. This is due to both a reduction in exploratory behaviour (resulting in more movement directly toward the reward) and a increase in speed due to ‘anticipation-invigoration.’

After D2 antagonist intervention the average speeds remained low throughout the course of the simulation. The small increase was due to less engagement in exploratory behaviour as learning in the core took place so that the agent perfectly targeted the marker however at slow speed.

In the next section we test robustness of the model on a real robot.

### 3.2 Real Robot

Fig. 4B shows the results from the real robot run. The x axis indicates again the individual trips along the runway to the food disk. Every trial the robot is put back at the starting position and then drives down the runway. After 72 trials the weights were reset and the whole experiment repeated 20 times. This resulted in a total of 1440 runs. The y axis is the average speed normalised to the maximum speed of the robot is able to achieve. The error bars result from the 10 runs for every trial. The upper graph (solid line, “No intervention”) shows the system in its normal operation: it learns to approach the food disk from the distance and it also increases its speed due to an increase in response vigour. This result confirms the findings from the simulation (Fig. 4A). As expected the real environment makes learning slightly more difficult resulting in a slightly slower convergence. This was due to the usual problems when dealing with real world scenarios such as robot and environmental imperfections, for example gearbox friction and a slightly uneven runway. Overall this shows that the model is robust in a real environment. An interesting difference is the that the highest standard deviation in speed is now before and during learning. This is due to the motors used which run at constant torque so that at low speeds the robot is affected more by the unevenness of the surface.

To test again, that the increase in speed is actually due to the increase in tonic DA and not just through the targeting of the food disk we again applied an D2 antagonist into the NAcc shell. Again, it can clearly be seen that the speed is much lower in the “D2 antagonist” scenario (dashed line) than in the “no intervention” scenario. Note that the speed increases in a similar amount than in the simulation but starts at a higher baseline to make sure that the robot moves all the time.

Overall the robot experiment confirms the results from the simulation.

### 3.3 Animal experiment vs robot experiment

In order to compare the data to that of the original experiment (Marx and Brownstein, 1963), the data was averaged over blocks of six trials to have a similar time course. A comparison of the averaged data and the data from the original experiment is shown in Figure 5. The average speeds for the simulation without intervention showed a similar rate and magnitude of increase to that of the original study. Note that the rats became about two times faster wereas with the robot we managed only about 25% increase which was mainly due to limitations of the motors of the robot. The dip in the average speeds of later trials in the original experiment was thought to be due to a satiation effect that carried over between days, this is not reflected in the simulation.

**Figure 5.**
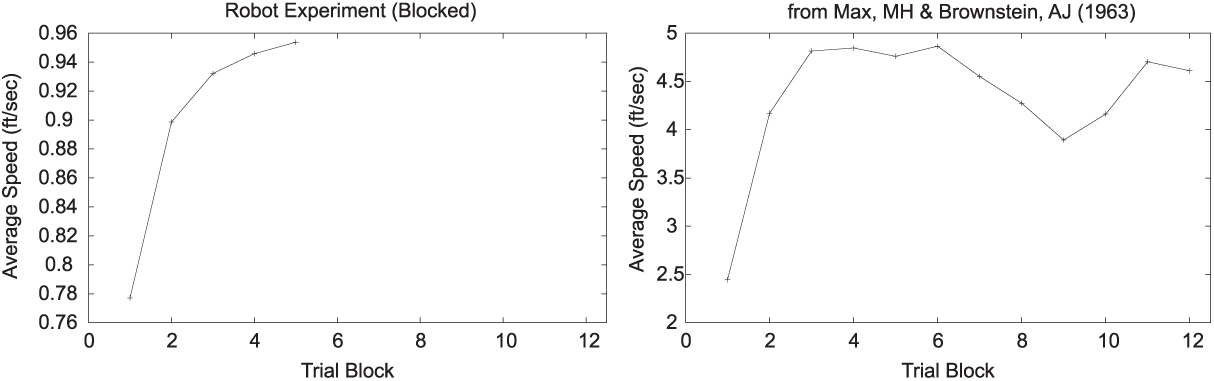
Comparison of the result of the robot experiment to the acquisition phase of the experiment described by Marx and Brownstein (1963). In order to make the experiments comparable we compressed the trials from the robot run (see Fig 4B) by merging 6 trials into one which we call “Trial block”. The y axis shows as before the average speed. The data in panel B is the original data from Marx and Brownstein (1963) replotted.

## 4 Discussion

In this paper we have presented a system wide circuit which is able to model response vigour. This has been achieved by integrating new anatomical connections projecting to/from the ventral striatum (Humphries and Prescott, 2010).

Tonic DA has long been implicated in motivational aspects of animal behaviour and has been reviewed extensively in Salamone et al. (2009); Wise (2004b). Dopamine antagonists consistently let animal work less for rewarding stimuli (Wise and Schwartz, 1981) and are less energised (Ikemoto and Panksepp, 1999). This has lead to the suggestion that tonic DA is responsible to energise actions (Salamone et al., 2009; Wise, 2004b). This is in accordance with our model where tonic DA boosts the motor output from the NAcc core due to the multiplicative effect tonic DA has on motor output. It is also easier to initiate an action with tonic DA present because it will be easier for the motor signal to reach the threshold to start a motor action (Salamone et al., 2009, 1995).

Formal theories for tonic DA have been recently developed around the reinforcement paradigm (McClure et al., 2003; Niv, 2007). It’s been stated that tonic DA represents the average reward value of the stimulus (or the context the animal is in) and that this value translates into response vigour. This is in accordance to our model where the average value is calculated in the NAcc shell (Dayan, 2001; Thompson et al., 2010b) and then translated into tonic DA via the shell-VP pathway. The circuitry around the shell has been extensively described in Thompson et al. (2010b) where phasic DA acts as an error signal and the shell calculates the average reward.

However we are at risk to become too dopamine centered in terms of motivation: the m-VP projects not just to the VTA where it causes a change in tonic DA but also to the MD thalamus which forms a loop with the limbic cortices (see Fig. 1,Zahm (2000)). This means that the shell disinhibits both the VTA and the MD thalamus in case of high activity in the shell. This pathway is therefore able to convey the reward value directly to the cortex and thus bypassing the dopaminergic system and consquently would be resistant against DA manipulations. It would be interesting to explore the differential effects of value transmission via the VTA and the MD.

The understanding of the anatomy of the wider NAcc circuitry has leaped forward in recent years by the discovery of the two subpopulations in the NAcc, namely the D1 receptor rich neurons and the D2 receptor rich neurons which has been reviewed in Humphries and Prescott (2010). Only a few year before this division was poorly understood and only menioned with caution (Wise, 2004a). Important in this context was the discovery of a direct and indirect pathway originating from these two subpopulations which made this model possible. Previous anatomical studies did not distinguish between these two pathways and thus obscured the action on the D2 receptors in the NAcc, for example (Zahm, 2000).

Knowing the circuitry around the value system and that dopamine is responsible to energise behaviour has implications for stimulation sites for DBS. While stimulation of the Nacc shell or VP will cause a change in tonic DA it will also change the activity in the MD thalamus. Weak stimulation of the VTA or MFB will just raise tonic DA but will not interfere with the MD modulation of the cortex. Knowing that tonic DA alone has already an energising effect might make MFB stimulation more attractive because it only affects DA concentration.

In this paper we have focused on the sub-cortical areas of the brain and that these compute the reward value or average reward, in particular the NAcc shell. However, there is growing evidence that the orbitofrontal cortex computes the reward value itself (Berthoud, 2004; Rolls, 1996, 2017) and then projects it to the NAcc shell which then in turn then controls the VTA. However, the underlying computations in the OFC are still poorly understood and further research needs to be undertaken to identify the different sub-nuclei which perform the distinct computations.

